# “Comprehensive Analysis of Nascent Transcriptome Reveals Diverse Transcriptional Profiles Across the Trypanosoma cruzi Genome Underlining the Regulatory Role of Genome Organization, Chromatin Status, and Cis-Acting Elements”

**DOI:** 10.1101/2024.04.16.589700

**Authors:** Pedro Leonardo Carvalho de Lima, Leticia de Sousa Lopes, Juliana Nunes Rosón, Alyssa Borges, Natalia Karla Bellini, Ana Tahira, Marcelo Santos da Silva, David Pires, Maria Carolina Elias, Julia Pinheiro Chagas da Cunha

## Abstract

Trypanosomatids are eukaryotic parasites exhibiting polycistronic transcription and trans-splicing. Post-transcriptional mechanisms are acknowledged as pivotal in gene expression regulation of their protein-coding genes. To comprehensively investigate the impact of transcription on gene expression in *Trypanosoma cruzi* and the association with the epigenetic landscape, we conducted a genome-wide nascent transcriptomic analysis. Our findings reveal significant asymmetrical transcriptional abundance across the genome, notably between polycistronic transcription units (PTUs) enriched in conserved genes (core PTUs) and those containing virulence genes (disruptive PTUs). We found that trypanosomes exploit linear genome organization to regulate transcription abundance by embedding virulence genes into highly transcribed core-enriched PTUs, by positioning PTUs near non-coding regions of small non-coding RNAs (e.g., tRNAs, snoRNAs), and by placing core CDSs in PTUs of various sizes. Additionally, we found correlations between open chromatin status and nascent transcript levels, both globally and particularly at transcription starting regions (divergent strand switch regions - dSSRs), indicating a crucial role for chromatin architecture in transcriptional regulation. While both core and disruptive dSSRs exhibit similar levels of some epigenetic marks (H2B.V deposition and 5mC), disruptive dSSRs display significantly higher 5hmC content and nucleosome occupancy compared to core dSSRs. Furthermore, we identified distinct conserved motifs within dSSRs of core and disruptive PTUs. These findings challenge the notion of constitutive and uniform transcription in *T. cruzi*, underscoring the paramount importance of linear genome organization, cis-acting motifs, and chromatin landscape in transcriptional regulation.

## Introduction

Gene expression is finely regulated by multiple mechanisms and layers, with transcription initiation standing out as one of the most tightly controlled processes. Chromatin-based mechanisms constitute the initial step, achieved by increasing its accessibility, followed by transcription itself, which involves multiple regulated steps such as the formation of the transcription preinitiation complex (PIC), initiation, promoter-proximal pausing, and pausing release (1). Both prokaryotic and eukaryotic genes typically contain promoter regions consisting of specific DNA sequence motifs that serve as platforms for the association of the transcription factors and transcription machinery, guiding them accurately to the transcription start sites (TSS). These promoters possess specialized sequences that have evolved to regulate gene transcription (2). Subsequent layers of gene expression regulation involve transcription elongation, termination, processing, and, for protein-coding genes, translation. All these layers can be finely regulated (1) and are crucial in determining the final levels of transcripts or proteins.

In trypanosomatids, protein-coding genes are organized in polycistronic transcription units (PTUs). It has long been accepted that genome and protein-coding genes are transcribed constitutively, with no regulation at transcription initiation sites (3–7). Evidence supporting the absence of transcriptional regulation in trypanosomatids existed even before the completion of genome sequencing in these organisms (3, 8–10). As a result, variations in mRNA abundances are mostly attributed to posttranscriptional mechanisms, of which differences in mRNA half-life play a significant role (7). Genome sequencing of trypanosomatids highlighted the presence of a few transcription factors, contrasting with the abundance of proteins containing RNA-binding motifs (11), likely associated with posttranscriptional regulation. Additionally, efforts to identify RNA Pol II promoters were elusive leading to the proposition that the “open” chromatin status could drive transcription initiation in the absence of promoters in any genomic region (9). Mathematical modeling of gene expression in procyclic forms of *Trypanosoma brucei* did not yield evidence for the regulation of RNA polymerase II transcription initiation. However, additional levels of regulation in the nucleus may play a key role in gene expression in other life forms (12). Collectively, these results indicate the lack of transcriptional regulation in these parasites. Consequently, significant effort was dedicated to elucidating the molecular components involved in posttranscriptional mechanisms in these organisms, which were recognized as excellent models for studying posttranscriptional regulation.

The genome sequencing of trypanosomatids also revealed the presence of divergent canonical histones, histone variants, and machinery associated with histone modification (11), which are known to be associated with gene expression regulation in many organisms (13). Thus, our group and others characterized an extensive atlas of histone post-translational modifications (PTMs) (14–20) and detected the presence of histone variants and PTMs, with base J marking transcription initiation and termination sites in trypanosomes (6, 21–24). In many organisms, including trypanosomes, genome organization and chromatin structure considerably impact gene expression, affecting transcription stability and localization. In this regard, we previously showed that open chromatin regions and a nucleosome-depleted region at 5’ UTR are associated with higher levels of mature mRNA (25, 26), indicating that chromatin reflects gene expression even in the context of polycistronic expression as in *T. cruzi*. Recently, three-dimensional genomic structures were also shown to be related to gene expression in trypanosomes (27, 28).

Although there is ample evidence that posttranscriptional mechanisms are critical for regulating mRNA abundances in trypanosomes (7), the current belief that trypanosomes do not regulate transcription initiation of RNA Pol II genes is being questioned. It has been known for decades that both *T. brucei* and *T. cruzi* exhibit a global decrease in transcription activity when replicative forms are compared to non-replicative forms, indicating that transcription regulation is evident during differentiation (8, 29, 30). Additionally, transcription of high-copy genes is lower than expected, suggesting that these genes may be subjected to transcription regulation in *T. cruzi* (29). More recently, GT-rich sequences and sequence-specific promoters were found to be necessary and sufficient to induce transcription initiation in dSSRs of *T. brucei* (31).

The *T. cruzi* genome was proposed to be compartmentalized into two main linear genomic regions: the “core compartment”, composed of conserved and hypothetical genes that are syntenic in other trypanosomatid species, and the “disruptive compartment”, composed of non-syntenic and repetitive multigene family (MF) genes, including the virulence factors trans-sialidase, mucins, and MASPs (32, 33). Recently, it was observed that these regions may fold differentially in the genome, with disruptive regions of *T. cruzi* showing fewer unprocessed transcripts than conserved regions (28). Extensive efforts to uncover the regulation of virulence factors (mainly the VSG family) expression in *T. brucei* reveal multifactorial players based on a combination of transcription regulation, DNA recombination, epigenetic mechanisms, and 3D organization (27, 34–36). In contrast to *T. brucei*, which has their virulence factors transcribed by RNA Pol I (37), the surface-coding virulence genes in *T. cruzi* are transcribed by RNA Pol II (8, 29). Thus, *T. cruzi* may use different strategies to ensure expression of their virulence factors. In fact, evidence of posttranscriptional regulation for individual sets of virulence factors (trans-sialidase, mucin, amastin) indicates that both infective and non-infective forms share similar transcription rates *in T. cruzi* (8, 38).

Aiming to comprehensively investigate whether all genomic regions in *T. cruzi* are consistently and evenly transcribed, we conducted a comparative analysis of PTUs containing conserved and virulence genes. This involved a thorough evaluation of nascent transcription expression alongside extensive data integration with epigenetic datasets and an assessment of conserved sequence motifs to elucidate our findings. Our results underscore significant differences in transcriptional expression primarily between core and disruptive regions likely driven by both cis (sequence and linear genome organization) and trans (chromatin/epigenetic modifications) factors, in addition to established post-transcriptional mechanisms.

## Materials and Methods

### Cell Culture

Epimastigote forms of *T. cruzi* strain Dm28c were cultured at 28°C in liver infusion tryptose (LIT) medium supplemented with 10% fetal bovine serum (FBS-Vitrocell), 0.4% glucose, 0.1 µM hemin, and 59 mg/L penicillin-G. Epimastigotes were maintained in the exponential growth phase by diluting them in fresh LIT medium to 2-4×10^6^ parasites/mL every 2-3 days.

### GRO-seq Assay

Parasite permeabilization followed established protocols (10, 29, 39). Briefly, 2 x 10^8^ epimastigotes were collected from cultures in the exponential growth phase (3200 rpm, 10 min, room temperature) in biological triplicates. The parasites were washed twice in transcription buffer (20 mM potassium glutamate, 3 mM MgCl_2_, 150 mM sucrose, 11.7 µg/mL leupeptin, 3 mM dithiothreitol). The pellet containing the parasites was resuspended in transcription buffer and chilled on ice for 5 min. Cell permeabilization was performed with 500 µg/mL Palmitoyl-L-lysophosphatidylcholine (Sigma-Aldrich) for 1 min at room temperature (RT), followed by washing with transcription buffer. Epimastigotes were then incubated (28 °C, 30 min) in run-on buffer (20 mM potassium glutamate, 3 mM MgCl2, 150 mM sucrose, 11.7 µg/mL leupeptin, 3 mM dithiothreitol, 0.6 mg/mL creatine kinase, 25 mM creatine phosphate, 400 U/mL RNAse Out, 1 mM ATP, 1 mM GTP, 1 mM CTP, 1mM UTP, 1 mM BrUTP). All experiments were conducted in biological triplicates (GRO_rep1, GRO_rep2, and GRO_rep3), and epimastigotes incubated in a run-on buffer without BrUTP were used as a background control sample (GRObckg). To confirm BrUTP incorporation into nascent RNA, a direct immunofluorescence assay was performed using Alexa Fluor anti-BrdU 488 antibody (Santa Cruz Biotechnology) and analyzed under an Olympus BX51 fluorescent microscope (100x oil-immersion objective) attached to an EXFO Xcite series 120Q lamp and a digital Olympus XM10 camera with Olympus Cell F camera controller software. Total RNA was extracted using Trizol® (Invitrogen) and treated with DNase I (Thermo Scientific) according to the manufacturer’s instructions. Immunoprecipitation of Br-UTP labeled RNA was performed using Dynabeads Protein G (Invitrogen) linked to Anti-BrdU antibody (Abcam). The magnetic beads preparation and RNA immunoprecipitation were performed according to the manufacturer’s instructions. Nascent RNA purification was conducted via ethanol precipitation (adapted from (40)). For this, 1 mL of cold 100% ethanol with 1 µg/µL glycogen and 300 mM NaCl were added to the samples followed by overnight incubation at −20 °C. Tubes were centrifuged (12000 x g, 30 min, 4 °C) and washed three times with 75% ethanol at RT (with 10 min incubation interval following each washing step) to dissolve the salts. Finally, the purified RNA pellet was air-dried, resuspended in 5 μL of nuclease-free water, and incubated for 15 minutes at RT. RNA quantification was performed with NanoDrop 2000 Spectrophotometer, and RNA integrity was evaluated through electrophoresis in a 1.5% agarose gel. RNA fragments (biological triplicates and a background sample) were deep sequenced using smRNA-Seq Kit for Illumina® (1 x 75 bp) at Polyomics Facility (Glasgow University), with no poly A selection or rRNA depletion. The GRO-seq dataset are deposited at PRJNA1073942.

### Bioinformatic Analyses

All experiments were run on a server configured with Ubuntu Linux Server 20.04, 224 cores, 1 TB of RAM, and 2 x Nvidia P40 GPU. The methods employed to analyze the GRO-seq data and public RNA-seq datasets are illustrated in S2 Fig and described below.

### Sequence Read Quality Check, Mapping, and Normalization

Raw sequencing files (*.fastq) were checked with FastQC version 0.11.9 (41). A quality filter was applied using Trimmomatic (42), which involved trimming adaptors, limiting read size to 75 bp, and removing low-quality bases (HEADCROP:3, LEADING:10, TRAILING:0, MAXINFO:65:0.05, MINLEN:35). Reads were then mapped against the *T. cruzi* Dm28c genome using Bowtie 2 v. 2.4.5 (43) (--verysensitive (-D 25 -R 4 -N 0 -L 19 -i S,1,0.40). When indicated, alignment files were filtered using a MAPQ (Mapping Quality) and sorted and indexed using samtools version 1.12 (2021) (Li, Handsaker et al. 2009). Data coverage was assessed using deeptools version 3.3.2 (Ramírez, Dündar et al. 2014), employing the BPM (bins per million) method within the bamCoverage function. Alternatively, Kallisto version 0.46.1 (44) was used to quantify each genomic feature in transcript per million (TPM) values using default parameters. An indexed transcriptome considering the entire genome separately by PTUs or CDSs was used (see below, S2B Fig). TPM values from experimental samples were subtracted by the corresponding feature found in the GRO background sample. RNA-seq datasets by Smircich et al., (2015) (45) (PRJNA260933) and Berna et al. (2019) (46) (PRJNA315252) were processed as described above.

### MNAse-seq and FAIRE-seq Analysis

Raw files were obtained from SRA under project numbers PRJNA763084 for FAIRE-seq (25) and PRJNA665060 for MNase-seq (26). Filtering and mapping were performed as described in the corresponding original papers.

### Extra Genomic Feature Annotation

Polycistronic regions, dSSRs, and cSSRs were annotated from the *T. cruzi* Dm28c GFF file downloaded from https://tritrypdb.org/ - DB53, Dm28c 2018) as previously described (25). To evaluate transcription along the entire genome, all coordinates with no annotation in the GFF file were further annotated as “others” using a custom Python v. 3.6.7 script available at https://github.com/PedroLeoardo/annotateOthers. If “others” were between CDSs (or between CDS and any ncDNA) or at the contigs edges, they were classified respectively as “other-IR (for intergenic regions)” or “other-end.” Core (conserved proteins and conserved hypothetical proteins), disruptive (mucin, MASP, and trans-sialidase), and both (GP63, RHS, DGF-1) CDSs were further annotated and visualized in a bed file. PTUs harboring 80% or more of CDSs from each genomic compartment were classified as “PTU core,” “PTU disruptive,” or “PTU both.” “PTU mix” contained all the remaining PTUs.

### PTU Abundance and DGE Analysis

Two strategies were used to evaluate if different PTUs would have similar transcription rates. TPM counts from PTUs considering the genomic coordinates of the entire PTU and the median of the TPM values from different CDSs of a given PTU were used. In all cases, TPM values were obtained from Kallisto version 0.46.1 and subtracted from the TPM counts from the background sample (GRO_bckg). DGE between infective (trypomastigote cells derived tissue cultures, TCTs) and non-infective (epimastigotes) forms was obtained from TriTrypDB (release 66) website considering DESeq2 analysis, fold changes of 2, p-adj<0.01 from a transcriptomic analysis (28).

### Data Visualization

Data coverage was normalized using the deepTools toolkit version 3.3.2 (47) by Bins Per Million (BPM) for GRO-seq and RNA-seq datasets, and Reads Per Genome Content (RPGC) for FAIRE-seq and MNase-seq datasets using the bamCoverage function to generate normalized coverage files (bigWig). Following normalization, landscape profiles and heatmaps from specific genomic features were systematically generated using the computeMatrix tool followed by plotHeatmap or plotProfile function from deeptools version 3.3.2. The Integrative Genomics Viewer (IGV) version 2.11.9 (2021) (48) was used for additional inspection and visualization.

### dSSRs Analysis

#### Motifs analysis

MEME platform version 5.5.5 (49) was used to identify conserved motifs in dSSR sequences. Only dSSRs between PTUs from the same genomic compartment (either core or disruptive) were considered. The motifs identified in both core and disruptive dSSRs were compared with known motifs in the TFBSshape database (50) using the TomTom platform version 5.5.5 (51). Transcription factors identified belonged to different phylogenetic groups.

#### Modified Bases Analysis

Modified base analysis was performed using public Oxford Nanopore sequencing datasets available at (28). Files were converted from FAST5 format to POD5 format using the pod5 library version 0.3.6 (pod5 convert fast5) (https://github.com/nanoporetech/pod5-file-format), which were used to call methylation (m) and hydroxymethylation (h) cytosines with the Dorado program, version 0.5.3 (dorado basecaller hac,5mCG_5hmCG) (https://github.com/nanoporetech/dorado). The readings were mapped using the minimap2 program (52) through the interface offered by the Dorado program itself against the genome of the Dm28c 2018 strain (version 60). The analysis of methylated bases was carried out using the modkit program, version 0.2.5 (https://github.com/nanoporetech/modkit). The modkit pileup subcommand generated files of the type bedMethyl (modkit pileup --interval-size 1000000) and BEDGRAPH (modkit pileup --bedgraph) on the positive or negative strands.

#### Epigenetic Enrichment Analysis

Using core and disruptive dSSRs coordinates, nucleosome enrichment was extracted by BAMscale version v1.0 (53) using BAM files from MNase-seq dataset (26) (FPKM values); H2B.V enrichment by H2B.V Chip-seq (ChIP/input) (24) using BigWigSummary from deeptools (47)) (average Raw Counts); 5mC and 5hmC enrichment obtained as described above were extracted from bedgraph files (column Nmod / Nvalid_cov).

### Statistical and downsampling analysis

Statistical were conducted using GraphPrism 8.4.3 or R 4.3.0. Downsampling involved randomly selecting 100 CDSs without replacement, repeated 50 times. Spearman correlation analyses were performed using the subset() function in R, followed by Spearman correlation tests using the cor.test() function for each genomic compartment, evaluating combinations such as “FAIRE vs RNA” and “FAIRE vs GRO.” The cor.test() function was iteratively executed 50 times to ensure robustness, generating stored p-values. To enhance accuracy, false discovery rate (FDR) correction was applied using the p.adjust() function. Spearman’s correlation was selected for its rank-based nature, making it appropriate for non-parametric data and providing robustness against outlier measures. Additionally, a customized R script was utilized to assess the significance of the data using the chi-square test of independence and Fisher’s test for calculating p-values. The distribution of features in the genome served as a control. Fisher’s exact test was employed for frequencies less than 5.

## Results

### Optimization of GRO-seq Assays in *T. cruzi* and Assessment of Data Quality

To evaluate transcriptional regulation and address whether the *T. cruzi* genome undergoes widespread and constant transcription, we conducted genome-wide sequencing of nascent transcripts in biological triplicates (GRO 1-3). A control group consisting of unlabeled parasites was utilized as a background sample (GRObckg). Figs S1A-B demonstrate efficient permeabilization of parasites, labeling with Br-UTP, and RNA extraction without degradation. Over 30 Mb reads were sequenced in all samples (S1 Fig, Table S1). To validate that our GRO-seq assay results were enriched in nascent transcripts, we established a bioinformatic analysis pipeline (S2 Fig), considering both coverage (in BPM) and TPM counts. To achieve this, counts from all genomic features, either classified or not by polycistronic regions (PTUs) (S2B Fig, Tables S2-3), were utilized. GRO-seq data was compared with public RNA-seq datasets (45, 54), serving as controls for nascent transcription expression as these datasets capture the abundance of mature transcripts resulting from a balance between their synthesis, processing (trans-splicing and polyadenylation), and degradation. S1C Fig illustrates that the distribution profile of read coverage in coding regions (CDS) from nascent (GRO-seq) and mature (RNA-seq) transcripts differs significantly. The presence of transcript processing (indicated by arrows) is evident when evaluating mature transcripts due to the depletion of reads upstream of the beginning of the CDS (ATG - S) and downstream of the end of the CDS (Stop codon – E). This pattern indicates mRNA processing and the formation of mature transcripts with the removal of upstream and downstream regions at the 5’ and 3’ UTR, respectively. In line with the absence of processing in nascent transcripts, this depletion is not observed in these regions in the GRO-seq data. The lack of transcript processing was detected regardless of the genomic compartments (as will be discussed below) (S1D Fig). The presence and absence of RNA processing were also evident by evaluating the TPM counts in CDSs and intergenic regions. We detected a log_2_ fold change of −0.16 between CDSs and their intergenic regions in nascent transcripts. In contrast, intergenic regions from mature transcripts exhibited a 1.7 log_2_ fold decrease compared to CDS regions (S1E Fig). The absence of processing was also evident at the spliced leader (SL) locus (S1F Fig), where the entire SL transcript of approximately 140 bp was detected in GRO-seq samples, while a processed 39 bp-SL transcript was detected in the mature transcriptome. Taken together, these results indicate that GRO-seq datasets are enriched in unprocessed RNA transcripts and, therefore, enriched in nascent transcripts.

### Distribution and Abundance of Nascent Transcripts Varies Across the *T. cruzi* Genome

We initially evaluated the transcriptional activity throughout the trypanosome genome, investigating whether specific genomic regions exhibited no preferential transcription, indicated by TPM values (GRO-GRO bckg) <= 0 (S3 Table). To achieve this, we considered all *T. cruzi* genomic regions (37,959 regions – S2C Fig) using a modified version of the annotation GFF file (see Star Methods section). We observed that 9% (4.7 Mb) of the genome (in bp) exhibited undetectable levels of nascent transcripts (Fig 1A). Among these regions, transcripts from non-coding DNA (ncDNAs), mainly snRNAs and snoRNAs, as well as from the contig ends, were overrepresented (chi-square test, p-value 10^-12^) (Fig 1B).

**Fig 1.**
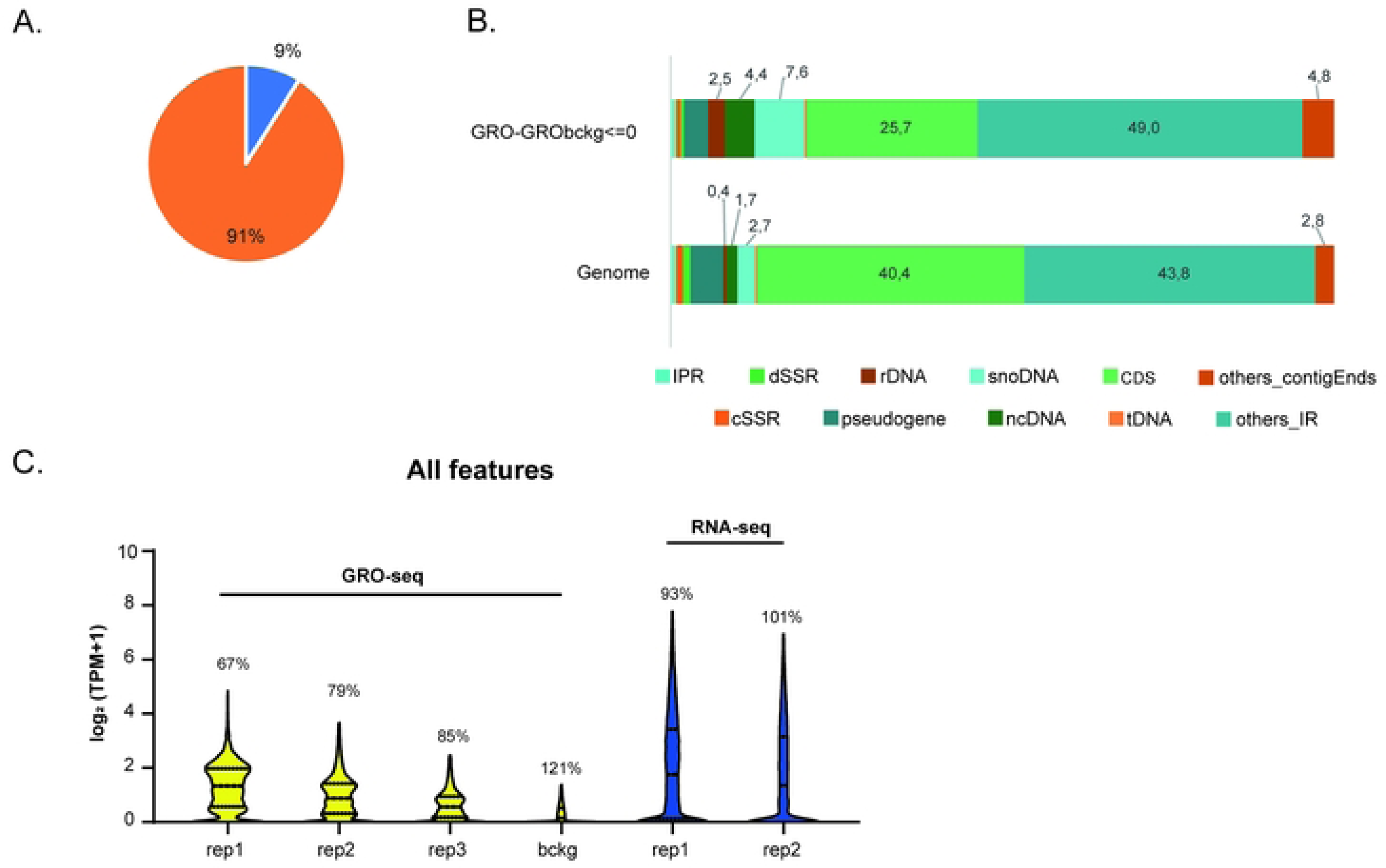
The Distribution and Abundance of Nascent Transcripts Across the *T. cruzi* Genome. **A** Percentage of *T. cruzi* Genomic Features Exhibiting No Nascent Transcription Expression (GROexp – GRO bckg ≤ 0). **B**. Bar plots representing the distribution of genomic features in GROexp – GRO bckg ≤ 0 (top) compared to the expected genome distribution (bottom). **C**. Distribution of Transcripts Per Million (TPM) Counts (log2(TPM +1)) across all Genomic Features (37,959 regions) from GRO and RNA-seq Assays. Coefficients of Variations are depicted above each violin plot. Outliers were removed (Q=1%).

We also detected an asymmetrical distribution among TPM counts of nascent transcripts (ranging from 0 to 7.8 in log2 (TPM+1)) (Fig 1C), indicating variability in nascent transcript abundance among different genomic features. The coefficient of variation (CV) was higher for mature transcripts (average 97%) than for nascent transcripts (average 77%), as expected due to intense post-transcriptional mechanisms in trypanosomes (7). However, the higher variation among transcript abundances in the nascent transcriptome indicates non-homogeneous transcription across the trypanosome genome, contrasting with previous thoughts.

### Assessment of nascent expression in PTUs revealed that core PTUs are more abundant than disruptive PTUs

The protein-coding genes of trypanosomes are organized into PTUs flanked by transcription start sites (TSS) and transcription termination sites (TTS) located at divergent and convergent strand-switch regions (dSSRs and cSSRs, respectively) (55). In *T. cruzi* (Dm28c strain), protein-coding genes are distributed among 1,286 PTUs. Polycistronic transcription precludes transcriptional control at the individual coding sequence (CDS) within the same PTU but does not preclude transcriptional differences among PTUs. To evaluate whether different PTUs share the same transcription rate, we compared their transcription abundance. As expected, TPM values from PTUs differ mostly within mature transcripts rather than nascent transcripts (Fig 2A). Nascent and mature transcript abundances from PTUs have coefficient of variation (CV) values of 66% and 72% (on average), respectively, implying that nascent transcript abundance is also not homogeneous among PTUs.

**Fig 2.**
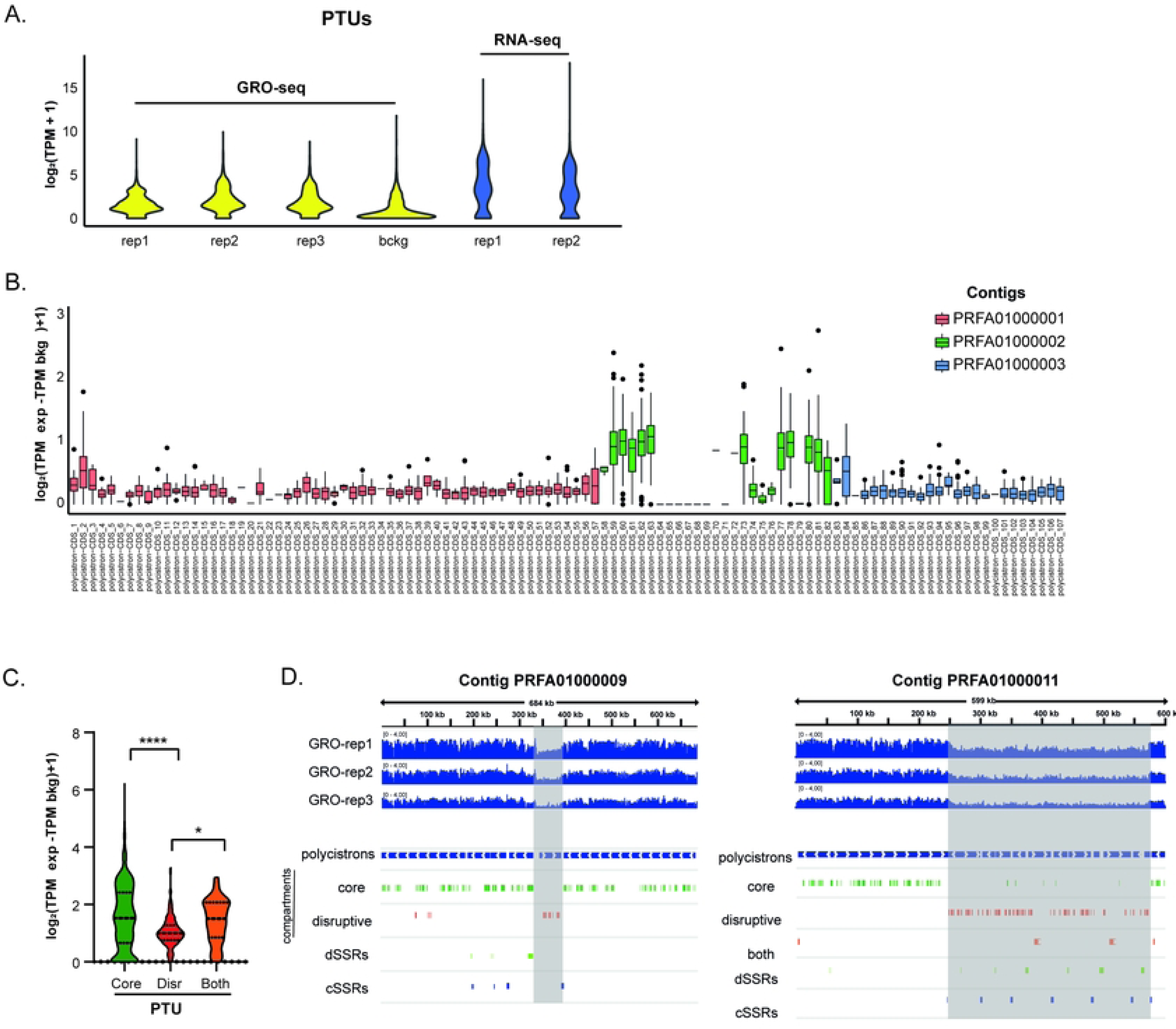
Assessment of Nascent Expression in PTUs Reveals Greater Abundance in Core PTUs Compared to Disruptive PTUs. **A** Distribution of TPM Counts (log2(GRO-GRObckg +1)) across all PTUs (1218) from GRO and RNA-seq assays. **B.** Boxplots of TPM Counts from PTUs on the first three major contigs (PRFA:01000001-01000003). Individual TPM Values (averaged among biological replicates) for each CDS are plotted for each PTUs. **C.** Violin plots of TPM Distribution from PTUs Containing 80% of Core, Disruptive, or Both Genes, respectively. **D.** IGV snapshot showing read coverage (BPM) from three replicates of GRO along two *T. cruzi* contigs (PRF01000009 and PRF010000011). Polycistrons are depicted in blue rectangles, while green and red rectangles indicate CDSs from Core and Disruptive compartments, respectively.

For a more detailed assessment, we evaluated the transcript abundance profile of the first hundred PTUs (Fig 2B and S4A Fig) distributed in the three major contigs (PRFA01000001, PRFA01000002, and PRFA01000003), highlighting that nascent expression profiles differ among PTUs. Interestingly, PRFA01000002 contains PTUs that are more abundant than those from the other contigs. This contig is enriched in CDSs of syntenic and conserved genes, which are part of the core compartment, whereas the lower abundant PTUs are mainly composed of non-syntenic/virulence genes that form the disruptive compartment.

To further evaluate whether these two compartments have different nascent expression profiles, PTUs composed of at least 80% of CDSs either from the core or disruptive or both compartments (composed of DGF-1, GP63, and RHS) were respectively classified as “core,” “disruptive,” or “both,” and their nascent expression profile was measured. Fig 2C-D and S3A Fig show that PTUs from the disruptive compartment indeed have lower nascent expression than those from the core and both compartments (One-way Ordinary ANOVA, p.adj<0.0001). Additionally, the nascent expression profile of individual CDSs from the core and disruptive compartments confirms that the former is more abundant than the latter (S3B Fig). To account for the repetitive nature of the disruptive compartment (33) and potential mapping issues, we used highly-quality mapped reads (MAPQ1 ≥ 30) (S3C Fig) and an RNA-seq dataset enriched in reads from disruptive regions (S3D Fig) to confirm the differential expression of these compartments. Taken together, these results indicate that virulence factor genes (disruptive compartment) in epimastigote forms have lower nascent transcription rates when compared to conserved genes (core compartment).

### Nascent Transcription of Core PTUs Depends on the Number of CDSs and Proximity to ncDNA Loci

To delve deeper into the differential transcriptional expression between core and disruptive compartments, we classified core and disruptive PTUs based on the number of CDSs per PTU. We observed that core PTUs containing more than 4 CDSs (369 PTUs) exhibited higher nascent expression compared to core PTUs with one (162 PTUs) or 2-4 CDSs (128 PTUs) (Fig 3A, S4A, and S4B, One-way ANOVA, p-adjusted <0.0001). Intriguingly, no correlation was observed between the number of CDSs per PTU within the disruptive compartment (Fig 3B, One-way ANOVA). Despite the median size (in bp) of core PTUs being larger than disruptive PTUs (S4C Fig), the length of PTUs increased with the number of CDSs, both for core and disruptive PTUs (S4C Fig). Hence, the lack of association between the number of CDSs in disruptive PTUs and nascent expression is unclear but aligns with the distinct nascent expression patterns observed in these two compartments.

**Fig 3.**
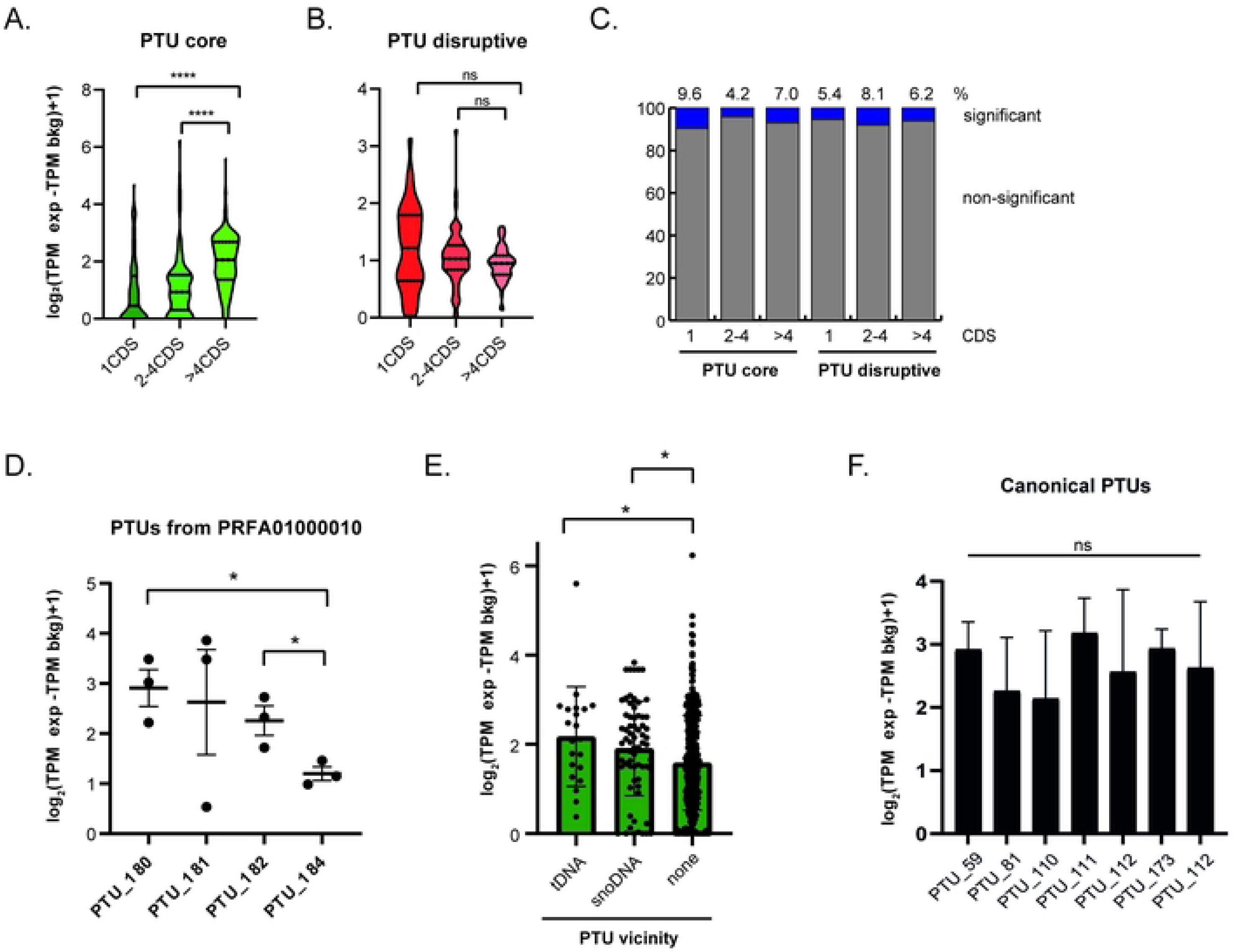
Nascent Transcription of Core PTUs Depends on the Number of CDSs and the Proximity of ncDNA Loci. **A** Violin plots illustrating the TPM distribution from PTUs containing 80% of core (left) or disruptive (right) genes, categorized by the number of CDSs per PTU. Core PTUs are divided into those with one (162 PTUs), 2-4 (128 PTUs), or 4 or more CDSs (369 PTUs). Disruptive PTUs are categorized similarly with one (40 PTUs), 2-4 (68 PTUs), or 4 or more CDSs (65 PTUs). Statistical analysis was performed using one-way ANOVA with Tukey’s range test at 95% confidence level (**** p<0.0001**). B.** Percentage of pairwise PTU comparisons that are statistically significant, classified by genomic compartments and CDS number per PTU using one-way ANOVA with Tukey’s range test at 95% confidence level. **C.** TPM counts from four core PTUs with 4 or more CDSs located on the same contig. **D.** Abundance values from PTUs not adjacent to any ncDNA (none) or adjacent to a tDNA or snoDNA locus. Statistical analysis was conducted using the Kruskal-Wallis test (* p<0.01). E. TPM counts of seven canonical PTUs (#59 - contig 2 - 100 CDSs, #81 - contig 2 - 56 CDSs, #110 - contig 4 - 129 CDSs, #111 - contig 4 - 74 CDSs, #112 - contig 4 - 59 CDSs, #173 - contig 9 - 19 CDSs, #181 - contig 10 - 57 CDSs). TPM values for each PTU were obtained considering the entire PTU coordinates. No statistical significance was observed.

Subsequently, we explored whether nascent expressions varied within PTUs previously classified based on their genomic compartment and number of CDSs. Between 5.4% to 8.1% and 4.2% to 9.6% of the pairwise comparisons were significantly different within disruptive and core PTUs, respectively (Fig 3C) (ANOVA test using Tukey’s range test as post-hoc, p <0.01). Among the differentially abundant core PTUs with four or more CDSs located on the same contig, we observed that differential expression was associated with PTUs positioned adjacent to a tRNA locus or at contig ends (Fig 3D). Among the top 20 differentially expressed PTUs, 35% contained ncDNA (mainly tDNAs and snoDNAs) adjacent either upstream or downstream. To assess whether the presence of ncDNA could globally interfere with transcriptional expression, we measured nascent expression from PTUs adjacent to tDNA and snoDNA compared to those without any associated ncDNA (Fig 3E). PTUs adjacent to tDNA and snoDNA exhibited, on average, higher transcriptional rates than those without any nearby ncDNA. This suggests that the presence of ncDNA may regulate *T. cruzi* PTU transcriptional expression.

Finally, we investigated whether nascent transcription levels indeed differed among “canonical core PTUs.” We manually selected seven PTUs that met specific criteria: i. contained more than 4 CDSs; ii. were located between intergenic regions delimited by dSSRs and cSSRs; iii. were not at the chromosome/contig end; iv. were not interrupted by tRNA, snoRNA, snRNA, or rRNA loci; and v. contained only genes from the core compartment (Fig 3F). No statistical differences were found among these PTUs, indicating that PTUs with similar features (related to genomic compartment, number of CDSs, and genomic location) exhibited the same transcription rate.

### Virulence factor genes located within core PTUs exhibit higher transcriptional rates than those located within disruptive PTUs

Since disruptive PTUs exhibit lower nascent expression than core PTUs, we investigated whether a disruptive CDS within a core PTU (classified as >80% of core CDSs) would show higher nascent transcription levels compared to those disruptive CDSs within disruptive PTUs (classified as >80% of disruptive CDSs), and vice versa for core CDSs within disruptive PTUs (Fig 4A). We found that core CDSs within disruptive PTUs had lower expression compared to core CDSs from core PTUs. Similarly, disruptive CDSs within core PTUs exhibited higher expression compared to those disruptive CDSs from disruptive PTUs (Fig 4B). Virulence factors such as trans-sialidases, mucins, and MASPs are part of the disruptive compartment (46). Therefore, we evaluated the nascent expression of these genes separately (Fig 4C). Consistently, we observed higher nascent expression levels for trans-sialidases, mucins, and MASPs when they were located within core PTUs compared to disruptive or mixed PTUs. Interestingly, genes for trans-sialidase type I, II, and VIII are overrepresented in core PTUs, while MASP and mucin are underrepresented (Fig 4D).

**Fig 4.**
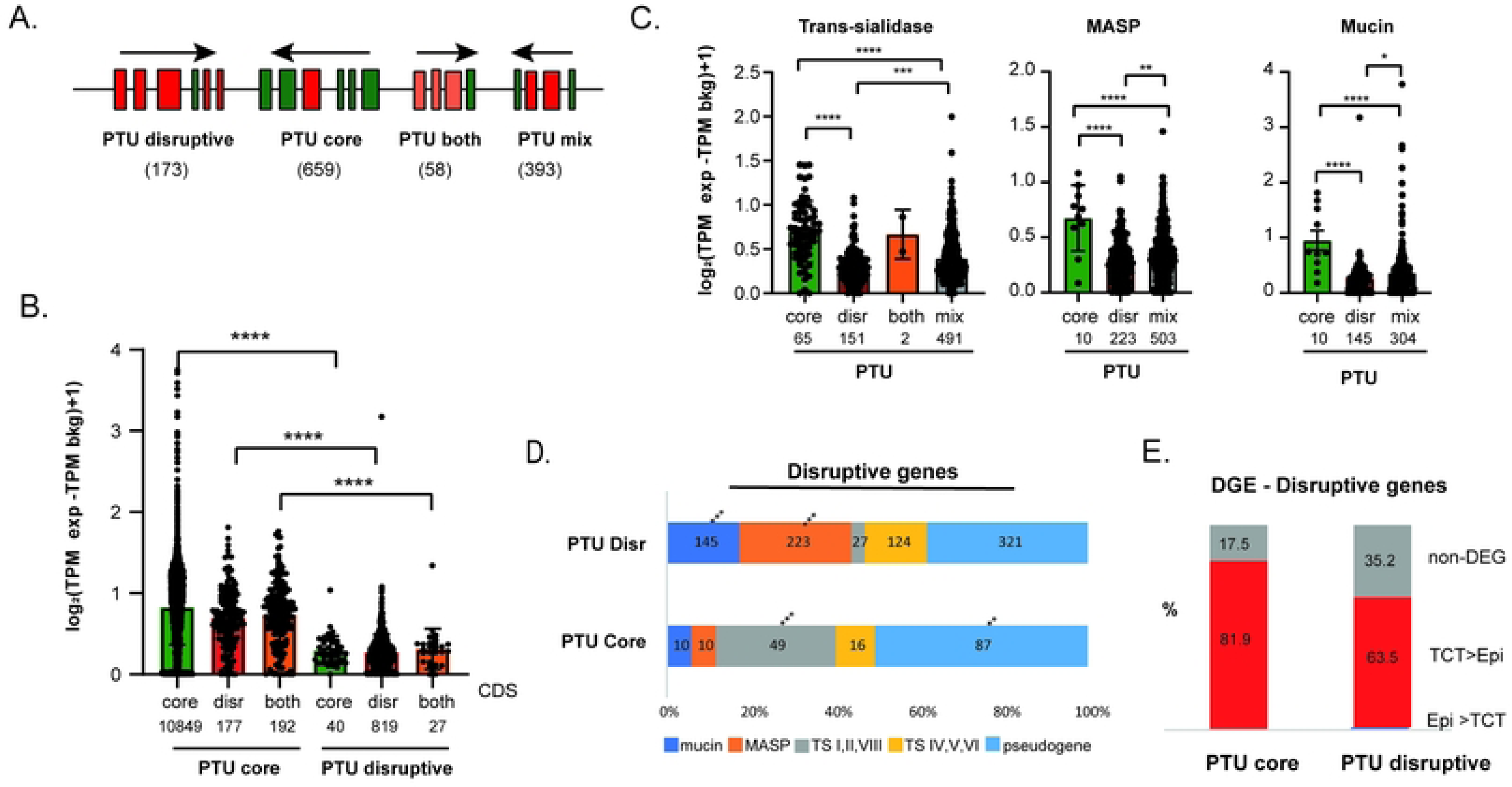
Nascent Transcription Expression Depends on the PTU Context. **A** Schematic representation of four PTU categories based on the content (at least 80%) of core (PTU core), disruptive (PTU disruptive), both (PTU both) genes, and mixed PTUs composed of core, disruptive, or both CDSs with less than 80% representation. The number of PTUs in each category is shown. Arrows indicate the transcription direction of PTUs. **B.** Bar plots of TPM counts from individual CDSs classified as core, disruptive, or both, from PTUs with more than 80% composition of core, disruptive, and both CDSs, respectively. **C.** Similar analysis shown in B for trans-sialidases, mucin, and MASP CDSs. Statistical analysis was conducted using one-way ANOVA with Tukey’s range test as post-hoc (p<0.0001). **D.** Distribution of trans-sialidase, mucin, MASP, and pseudogenes in core and disruptive PTUs (chi-square test with a p-adjusted < 0.0001). ** for adjusted p-value < 0.001, *** for adjusted p-value < 0.0001 (chi-square test). **E.** Percentage of differentially expressed disruptive genes (Epimastigotes versus TCTs, p-adjusted <0.01, fold change of 2, based on (28)) located in core or disruptive PTUs.

It is known that *T. cruzi* infective forms contain higher levels of mature transcripts from virulence factors (54); however, these levels are regulated post-transcriptionally since infective forms do not increase transcription levels of virulence factors compared to non-infective forms (8, 10, 38). Here, we demonstrate that virulence-factor genes are more transcribed when located within core PTUs. Thus, we hypothesize that genes preferentially enriched in infective forms (cellular trypomastigotes) originate from those located preferentially in core PTUs. Accordingly, we found that 82% of virulence-factor genes located in core PTUs are upregulated in TCT forms compared to 63% of those located in disruptive PTUs (p-adj <0.01, fold change of 2, Fig 4E). These findings suggest that genome organization, particularly linear gene order, plays a critical role in regulating transcription to ensure the expression of essential virulence-factor genes in *T. cruzi*.

### Open chromatin status, both globally and locally at dSSRs, correlates with nascent transcript levels

Regions with more open chromatin are generally associated with higher transcriptional activity in several eukaryotes (56). Previously, we demonstrated a positive correlation between open chromatin (based on FAIRE-seq data) and nucleosome occupancy (measured by MNAse-seq data - depth of the nucleosome-depleted region at 5’UTR) with mature transcript levels (25, 26). Here, we show a stronger correlation between nascent transcriptome and open chromatin status (ρ = 0.65) than mature transcriptome and open chromatin (ρ = 0.43) (Fig 5A). Furthermore, we classified genomic regions into three groups (high, medium, and low) based on levels of open chromatin (measured by RPGC) and compared them with nascent expression levels. As expected, higher levels of open chromatin corresponded to higher nascent expression (Fig 5B).

**Fig 5.**
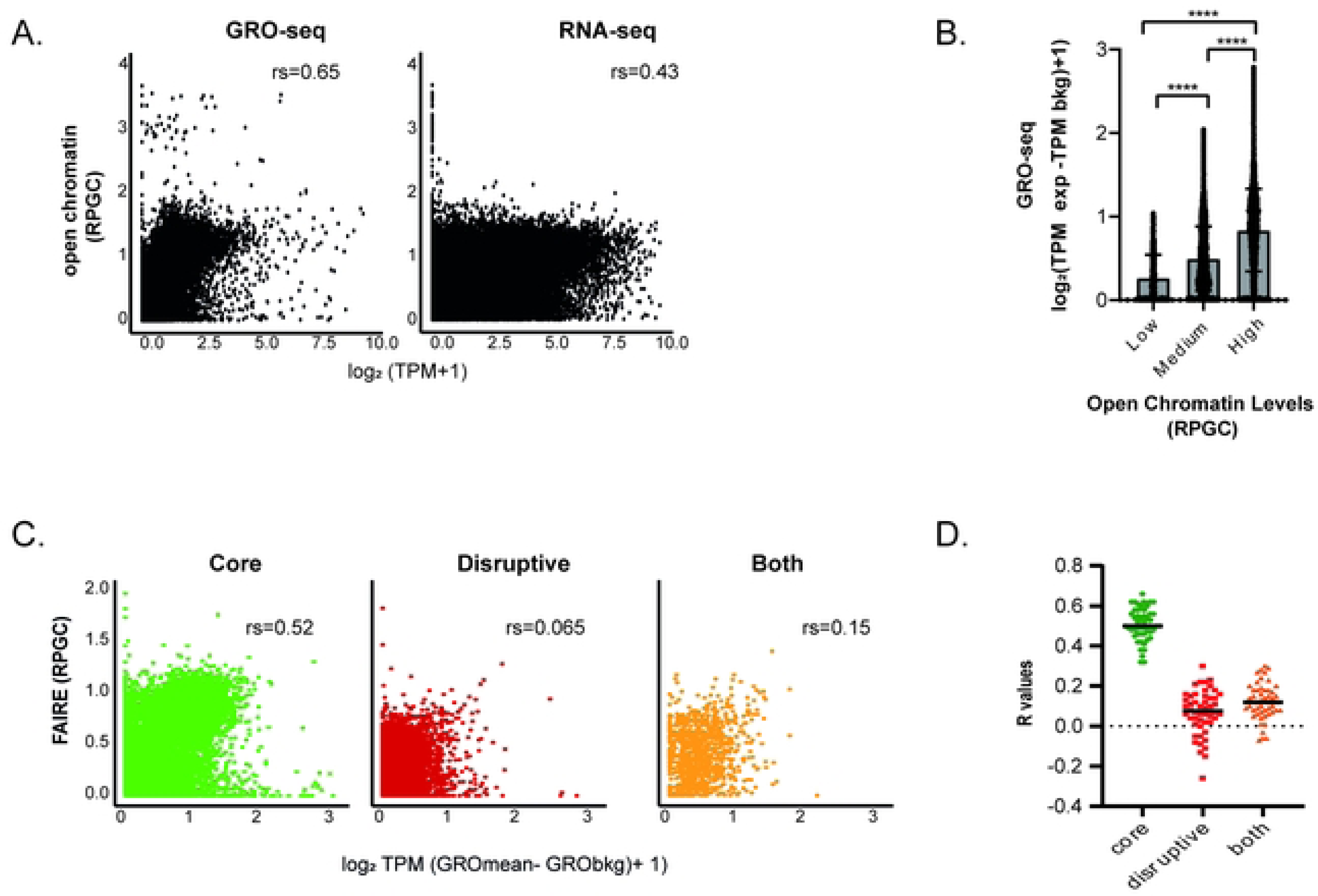
Global Open Chromatin Status at Core Region Correlates with Nascent Transcript Levels. **A** Scatter plots of open chromatin (FAIRE-seq dataset, RPGC levels) profiles versus nascent (GRO-seq) or mature (RNA-seq) transcriptomes for all *T. cruzi* genomic features. R values are indicated. **B.** Bar plots illustrating TPM counts of genomic regions based on their open chromatin levels (high - 25% highest RPGC levels, low - 25% lowest RPGC levels, medium – the 50% remaining). Outliers were removed using the ROUT method, Q=1% at PRISMA. **C.** Scatter plots of open chromatin (FAIRE-seq dataset, RPGC levels) profiles versus nascent (GRO-seq) transcripts for coding DNA regions classified as core, disruptive, and both regions. R values are displayed. **D.** R values of open chromatin profile and nascent transcription obtained through downsampling analysis of a randomly selected 100 genes (for the indicated compartment) repeated 50 times. Statistical analysis was performed using one-way ANOVA with Tukey’s range test as post-hoc (****p<0.0001).

Remarkably, we detected a moderate positive correlation (ρ = 0.52) between “open” chromatin and nascent transcription exclusively for genes from the core compartment, confirmed by down-sampling analysis (100 genes representing each compartment randomly selected 50 times) (Fig 5C). In contrast, for global analysis, no correlation was observed between nucleosome occupancy and nascent transcription, regardless of the genomic compartment (S5A-B Fig). These findings suggest that expression in core and disruptive compartments is subject to different transcriptional regulatory effects based on chromatin landscape.

Although the observation of a global association between chromatin landscape and nascent expression is intriguing for these organisms, establishing whether chromatin genuinely could influence transcription necessitates the examination of local variations, especially at transcription start regions. In trypanosomes, transcription initiates mainly at dSSRs (6, 57), which are known to harbor enrichment of open chromatin, H2A.Z/H2B.V, and active histone marks (6, 21, 24, 58, 59). Here, we demonstrate that in epimastigote forms, dSSRs can be clustered into three classes based on their levels of chromatin opening (Fig 6A). For simplicity, we termed dSSRs associated with cluster 1 as “open” and those with cluster 3 as “closed.” We observed that PTUs associated with open dSSRs have higher abundance of nascent transcripts than those associated with medium and closed dSSRs (Fig 6A).

**Fig 6.**
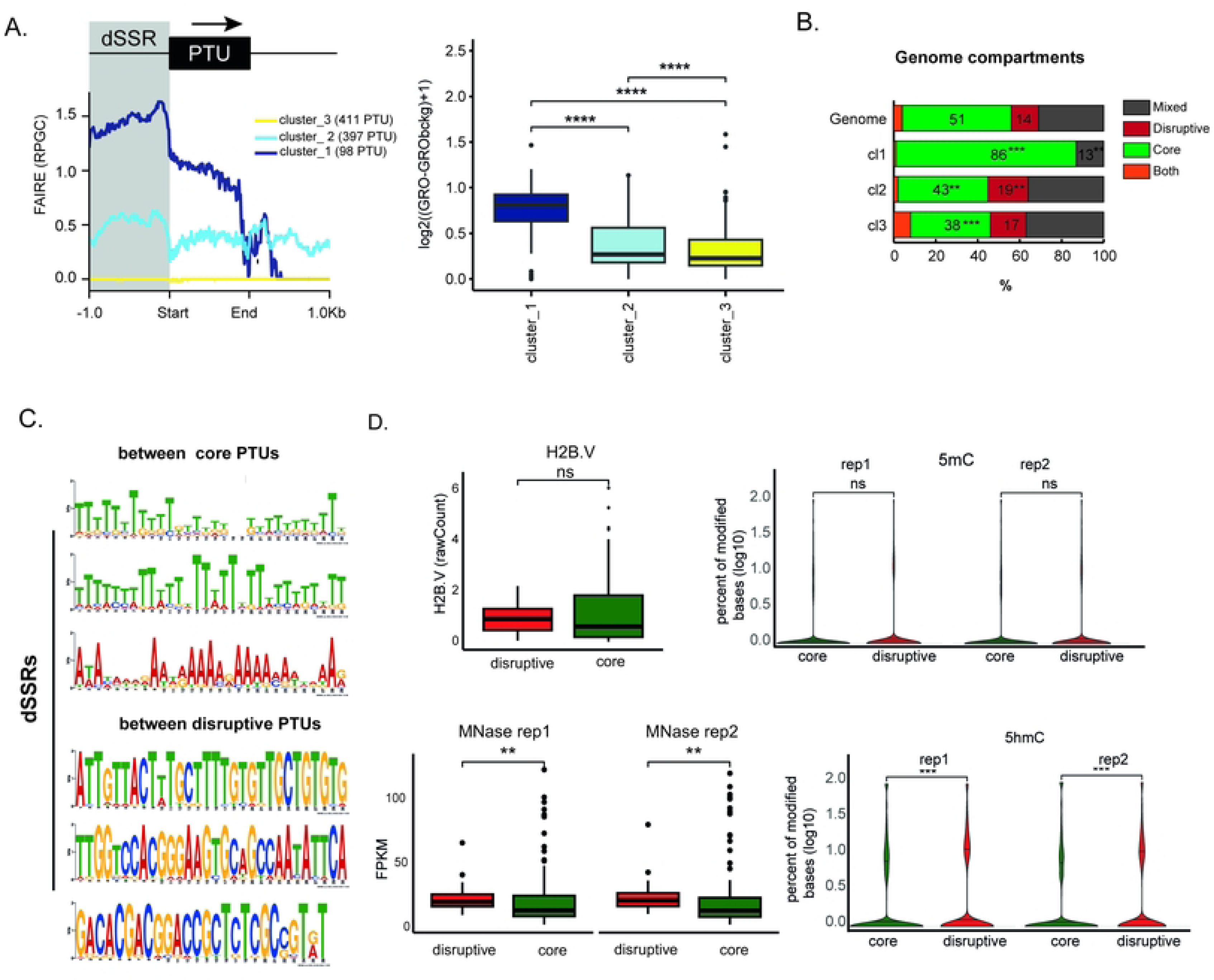
Core and Disruptive dSSRs Differ in Their Epigenetic Landscape and Sequence Motifs. **A** Left: Hierarchical cluster analysis of the distribution of the RPGC log2 ratio in PTUs from epimastigotes considering 1 Kb upstream or downstream, highlighting the dSSRs. Right: Bar plots depicting TPM levels from PTUs associated with dSSRs that harbor different levels of open chromatin as shown on the left. PTU abundance was based on median TPM values from CDSs of each PTU. Wilcoxon test, p.adj <0.05**. B.** Distribution of core, disruptive, both, and mixed PTUs associated with open and closed dSSRs as depicted in A. (chi-square test under a p-adjusted < 0.01). * for adjusted p- value < 0.0125, ** for adjusted p-value < 0.00125, *** for adjusted p-value < 0.000125 (chi-square test). **C.** Sequence motifs determined by the MEME platform using dSSRs from either core or disruptive PTUs. Only dSSRs between PTUs from the same compartment were analysed. The top 3 sequence motifs are shown. E-values were higher than 1.5e^-076^ **D.** Distribution of abundance levels for 5hmC and 5mC (Nmod/Nvalid_cov, bedgraph score of coverage), H2.BV (ChIP/input) deposition (average raw counts), and nucleosome levels (MNase-seq) (FPKM) for core and disruptive dSSRs. Mann-Whitney test, ** p.adj <0.01. Only dSSRs between PTUs from the same compartment were used in C and D analysis.

To gain further insights into these dSSRs clusters, we evaluated their size in base pairs, PTU compartment classification, and number of CDSs per PTU. Open dSSRs (cluster 1) are shorter (median size ∼1000 bp) than those with closed chromatin states (clusters 2 and 3) (S6 Fig). Comparing open (cluster 1) and closed (cluster 3) chromatin dSSRs with the expected genome distribution of core and disruptive PTUs, we found that the former has no disruptive PTUs while the latter has 17% of disruptive PTUs. Additionally, core PTUs are enriched (86%, chi-square p < 0.05) in open dSSRs and impoverished (38% of their PTUs, chi-square p < 0.05) in closed dSSRs (Fig 6B). Hence, it is evident that the disruptive compartment exhibits not only reduced open chromatin levels but also a higher prevalence of closed dSSRs. These findings strongly indicate that chromatin accessibility at transcription start regions correlates with the abundance of nascent transcripts, potentially influencing transcription rates. This offers a compelling molecular rationale for the regulation of transcription initiation.

### Transcription start regions from core and disruptive PTUs share similar levels of some epigenetic marks but differ in their nucleosome content, 5hmC levels and sequence motifs

Here, we demonstrated that PTUs composed of genes from core and disruptive genomic compartments exhibit distinct levels of transcriptional expression, which correlate with varying degrees of open chromatin at their dSSRs. Motivated by these findings, we explored whether core and disruptive dSSRs also exhibit differential levels of other epigenetic marks, including H2B.V deposition (a hallmark of dSSRs in trypanosomes), 5-methylcytosine (5mC), 5-hydroxymethylcytosine (5hmC), and nucleosome content. We found that while both core and disruptive dSSRs harbor similar levels of H2B.V and 5mC; disruptive dSSRs exhibit a higher 5hmC content and nucleosome density compared to core dSSRs (Table S5-S6).

Finally, we investigated whether dSSRs situated between core or disruptive PTUs would display differential conserved motifs that resemble putative promoter sequences, as recently identified in trypanosomes. Remarkably, we observed substantial differences in the enriched motifs within dSSRs from core versus disruptive PTUs. While the former exhibited enrichment of poly-T and poly-A repeats, the latter displayed a diverse array of motifs with varying nucleotide compositions, albeit with a predominance of G and T nucleotides (Fig 6C). Collectively, these findings underscore the divergent regulatory mechanisms based on cis (sequence) and trans (chromatin) features of core and disruptive dSSRs, suggesting a potential regulatory mechanism governing transcription initiation in *T. cruzi*.

## Discussion

Traditional transcriptome analyses are typically based on detecting the total set of mature transcripts, encompassing both unprocessed and processed RNA forms. To gain a deeper understanding of transcriptional activity, strategies such as “run-on” assays are employed. Such assays utilize labeled nucleotides during *in vitro* transcription coupled with large-scale sequencing techniques, such as Global Run-On sequencing (GRO-seq) (1, 60). This approach enables the detection of nascent RNA levels, providing a more precise understanding of global transcriptional regulation. By conducting a genome-wide analysis of nascent transcripts in *T. cruzi*, we identified an asymmetrical distribution of transcription abundance, indicating that genomic regions are not uniformly transcribed. Specifically, we observed different transcriptional levels between core and disruptive regions, with PTUs from disruptive regions being less transcribed than those from core regions. Furthermore, within core PTUs, variations in nascent expression levels were detected corresponding to the number of coding sequences (CDSs). Although the underlying reasons for these observations remain unclear, it has been previously demonstrated that longer PTUs exhibit anomalous expression in trypanosomes (61). It is important to note that among core PTUs with the same number of CDSs, no transcriptional differences were observed, indicating conserved levels of transcription among core PTUs of similar sizes.

In PTUs from the same genomic compartment and with a similar number of CDSs, few statistical differences were detected (less than 10%). These differences were mainly associated with the presence of *loci* containing snoRNAs and tRNAs in their vicinity. This finding suggests that the presence of non-coding RNA *loci* may interfere with adjacent transcriptional expression. Although the underlying mechanisms for that are unknown, we envisage that it may be associated with the fact that tRNAs and some snoRNAs are transcribed by a different RNA Polymerase, the RNA polymerase III (62). Basal transcripts factors of RNA polymerase III and II can be shared resulting in competition, which could result in transcription interference (63, 64). Additionally, some tDNA *loci* function as insulators, contributing to the 3D architectural organization of the genome, potentially impacting gene expression (65). However, this is yet to be proven in trypanosomes. Together, these data suggest that the trypanosome genome is organized to promote transcriptional gene regulation by positioning tDNAs and snoRNAs close to some protein-coding genes.

In *T. brucei*, virulence factors (VSGs) are transcribed unusually by RNA Pol I (37), and considerable effort has been dedicated to understanding the underlying mechanism of their gene expression (35, 66). In contrast, *T. cruzi* surface-coding virulent genes are transcribed by RNA Pol II, and efforts to uncover their regulation have found solid evidence of posttranscriptional regulation in a particular set of genes (8, 10, 29, 38). In this way, understanding how parasites regulate RNA polymerase II transcription of protein-coding genes may provide important insights into how *T. cruzi* regulates the expression of their repertoire of virulence genes. By evaluating the nascent transcriptome and linear organization of the *T. cruzi* genome, we observed a group of virulence genes embedded in core PTUs, exhibiting higher transcription rates. Notably, a significant proportion of these highly transcribed virulence genes are enriched in infective forms, suggesting a strategic genomic positioning to ensure robust expression. This finding reinforces the notion that the trypanosome genome is strategically organized to regulate gene expression. Linear genome organization has been shown to be critical in gene expression regulation in *T. brucei* (67). In these organisms, genes differentially expressed during the heat shock response are located either proximal or distal to transcription start sites.

Different organisms regulate gene expression by modulating the chromatin landscape. Consistent with a role of chromatin architecture in transcription regulation (68), we detected a positive correlation between open chromatin levels and nascent transcription. The coding regions of virulence factors (disruptive regions) are enriched in nucleosomes, depleted in open chromatin (25, 26), and form more inter-chromosomal contacts (28), suggesting that they are enriched in condensed and less accessible chromatin. In contrast, core regions are more open with fewer nucleosomes and exhibit higher transcription levels. Therefore, this chromatin landscape may be involved in modulating the binding of RNA polymerase II machinery and basal transcription factors, justifying the differential transcription rates in these regions.

Moreover, while a clear correlation between core nascent transcription and chromatin openness was observed, this correlation is not evident when analyzing disruptive regions. The reasons for that are unclear but suggest that virulence and conserved regions are under different regulatory players. While a clear correlation between core nascent transcription and chromatin openness was observed, such a relationship is not evident for the disruptive regions. Interestingly, levels of open chromatin and nucleosome abundance are not altered in disruptive virulence regions (compared to core regions) in infective forms when compared to non-infective ones (25), suggesting that the chromatin landscape of disruptive regions may remain silent and impervious to transcriptional regulation. Run-on experiments indicated a similar transcription rate of virulence genes in infective and non-infective forms, highlighting that trypanosomes employ tight post-transcriptional control to ensure high expression of virulence factor genes in infective forms (8, 10, 38). Thus, we hypothesize that disruptive genomic regions are silenced both in infective and non-infective forms, and parasites may take advantage of the presence of virulence factors located within core PTUs. Future experimental analyses should be conducted to verify the validity of this hypothesis.

While observing a global association between chromatin landscape and gene expression is intriguing, determining whether chromatin indeed impacts transcription requires examining local differential alterations, particularly at transcription start regions. In this regard, our previous observations revealed varying levels of open chromatin and nucleosomes in dSSRs between infective and non-infective forms (25). Focusing on non-infective forms in this study, we found that PTUs associated with open dSSRs exhibit higher transcription rates compared to those with closed dSSRs. This data indicates that open chromatin at regions specifically associated with transcription initiation may indeed play a critical role in gene expression, suggesting a mechanism of regulation at transcription initiation in *T. cruzi* that contrasts with previous proposals in the field.

Until recently, the only promoter sequence described for RNA polymerase II was found at spliced leader RNA genes in trypanosomatids (69). However, the discovery of GT-rich elements and a 75 bp promoter in *T. brucei* dSSRs challenged the previous belief that there were no promoter sequences for mediating RNA polymerase II transcription of protein-coding genes (6, 31). Here, we determined that transcription start regions from core and disruptive PTUs differ greatly in their nucleotide content. Disruptive dSSRs have a higher content of G and T, with some repeats of GT, as seen in *T. brucei* (6). Guanine-rich motifs are common features of promoter regions (70). On the contrary, dSSRs from core PTUs are enriched in poly-A and polyT sequences. Poly dA-dT elements, ranging from 15 to 30 base pairs, are commonly found in the genomes of all eukaryotes, particularly in promoter regions. Studies have demonstrated the importance of these tracts in transcriptional regulation and recombination (71, 72). Regions containing homopolymeric elements (dA-dT) form more rigid and less flexible structures than conventional DNA, enabling them to resist nucleosome entanglement and providing access sites for transcription factors (73, 74). Consistent with this, dSSRs from core PTUs harbor fewer nucleosomes than those from disruptive PTUs. Collectively, these data reinforce the idea that sequence-specific elements within core and disruptive dSSRs may determine whether a region becomes enriched in nucleosomes, suggesting a putative mechanism of transcription initiation regulation. If nucleosome content is primarily determined by these cis-elements, variations among life forms should not be expected. However, we found that dSSRs from infective forms are enriched in nucleosomes compared to non-infective (26). Thus, other factors may guide the deposition of nucleosomes in these regions, contributing to regulation. In this context, we found no association between the deposition of the epigenetic marks H2B.V and 5mC in core versus disruptive regions, but an increase in 5hmC in disruptive dSSRs. 5mC and 5hmC exhibit distinct distribution patterns in the genome of eukaryotes; while the former is more closely linked to gene repression, the latter is predominantly found in active enhancers and surrounding expressed genes (75). This may appear contradictory to our findings, as we detected lower expression associated with disruptive regions; however, it is still necessary to investigate the role of these cytosine modifications in trypanosomes to determine if they play the same role described in other eukaryotes.

In summary, this study challenges the paradigm that the genome of *T. cruzi* is constitutively and uniformly transcribed. Moreover, we demonstrate that even in genomes organized into PTUs it is possible to implement strategies to regulate transcription, such as positioning of virulent factor genes within higher transcriptional core PTUs or flanking PTUs with ncDNA *loci*. The wealth of biological resources is vast, and studying eukaryotes from different domains of life continues to provide surprises.

## DATA AVAILABILITY

GRO-seq, MNAse-seq, FAIRE-seq data can be accessed at the Sequence Read Archive (SRA) (https://www.ncbi.nlm.nih.gov/sra) with the following accession number: PRJNA1073942, PRJNA665060 and PRJNA763084. Oxford Nanopore sequencing datasets available at (28).

## AUTHOR CONTRIBUTIONS

Conceptualization: MCE and JPCC Data curation: PLCL, DP, JPCC

Formal Analysis PCL, LL, JNR, AB, NKB, AT, MSS, DP, JPCC

Funding acquisition MCE and JPCC

Investigation PCL, LL, JNR, AB, NKB, AT, MSS, DP, JPCC

Methodology PCL, LL, JNR, AB, NKB, AT, MSS, DP, JPCC

Project administration MCE and JPCC

Resources: MCE and JPCC

Software: PCL, DP, AT Supervision: JPCC, DP, AT

Writing – original draft: PCL, JPCC

Writing – review & editing: PCL, LL, JNR, AB, NKB, AT, MSS, DP, MCE, JPCC

## ACKNOWLEDGEMENTS

We thank Ivan Novaski Avino for technical assistance and Dr. Alex Ranieri Lima for important inputs on the data and bioinformatic analysis. We thank Herbert Guimarães de Sousa Silva, Dr. Ana Paula de Jesus Menezes and Dr. Simone Calderano for reading this manuscript and providing critical comments. We are also grateful to Dr. Sergio Verjovski de Almeida and Dr. Ariel Silber for critical suggestions.

## CONFLICT OF INTEREST

The authors declare no competing interests.

### Declaration of generative AI and AI-assisted technologies in the writing process

During the preparation of this work the author(s) used https://chat.openai.com/ to correct grammar, spelling, typos and improve readability and language. After using this tool/service, the author(s) reviewed and edited the content as needed and take(s) full responsibility for the content of the publication.

## Supporting Information

### Supporting Legends

**S1 Fig. GRO-seq Assays: Comparison of Nascent and Mature Transcriptomes. A.** Fluorescence microscopy images showing nascent RNA labeled with Br-UTP in the nuclear space of *T. cruzi* epimastigote forms. **B.** Agarose gel displaying RNA integrity extracted from parasites subjected to transcription assays with or without Br-UTP labeling. **C.** Summary plots of read coverage (median values) from coding DNA sequence (CDS) regions of three biological replicates (rep1, rep2, and rep3) of GRO-seq (top) (GRO/GRO bckg) and RNA-seq (bottom) ^45^, assays of *T. cruzi* epimastigote forms. Median coverage values were scaled individually for each replicate. **D.** Read coverage of three biological replicates of GRO-seq (left) (GRO/GRO bckg) and RNA-seq (right) assays of core and disruptive genomic compartments. **E.** TPM counts from CDSs and intergenic regions (IR) of GRO and RNA-seq assays. Median values are depicted above each boxplot. Median coverage values were scaled individually for each replicate. **F.** Summary plots of read coverage (median values) from the spliced leader locus of one representative GRO and RNA-seq sample. Yellow lines and boxes represent results from GRO-seq assays; blue lines and boxes from RNA-seq assays. The start (S – first ATG) and end (E – stop codon) of CDS are shown, along with 1 Kb upstream and downstream.

**S2 Fig.A.** Scheme of the bioinformatic pipeline used to retrieve coverage and TPM counts from GRO and RNA-seq datasets. B. Scheme of two GFF files obtained to cover all genomic features, considering either the genome classified by PTUs or by all CDSs. C. Distribution of genomic features considering GFF files from all features (37,959 regions) in the *T. cruzi* genome (Dm28c strain). IR stands for Intergenic Region; IPR - Intergenic polycistronic region is located between a polycistron and a locus of non-coding RNA.

**S3 Fig. A**. Top: Boxplots of TPM counts from PTUs of the first three major contigs (PRFA:01000001-01000003). Individual CDS TPM values for each biological replicate (GRO-seq) are plotted for each PTU. **B**. Left: Violin plot of the TPM distribution from individual CDSs of the core (conserved proteins and conserved hypothetical proteins), disruptive (mucin, MASP, and trans-sialidase), and both (GP63, RHS, DGF-1) compartments. Right: Confidence interval graph from the one-way ANOVA with Tukey’s range test as post-hoc (95% confidence). **** p<0.0001. **C**. Distribution of TPM values in core, disruptive, and both compartments based on MAPQ filters. D. IGV snapshot of GRO and RNA-seq coverages showing that disruptive regions (ref rectangules) are covered by transcripts in trypomastigote forms (TRYPO).

**S4 Fig. A.** Confidence interval graph of the one-way ANOVA with Tukey post-hoc (95% confidence) from Figure 3A. B. Violin plots of TPM distribution from PTUs containing 80% of core genes classified by the number of CDS per PTU. PTUs TPM values were based on the median TPM values from individual CDSs from each core PTUs with one (162 PTUs), 2-4 (128 PTUs), or 4 or more CDSs (369 PTUs). PTU length in bp classified by genomic compartment (C) and by compartment and number of CDS (D).

**S5 Fig. A.** Scatter plots illustrating the nucleosome occupancy (MNase-seq dataset, RPGC levels) profile versus the nascent (GRO-seq) transcriptome for coding DNA regions classified as core, disruptive, and both regions. R values are presented. B. R values of the open chromatin profile and nascent transcription obtained through downsampling analysis of a randomly selected 100 genes (for the indicated compartment) repeated 50 times.

**S6 Fig. dSSRs size distribution according to chromatin openness.** The first 20 contigs of *T. cruzi* strain Dm28c were examined for length of dSSRs located between PTUs within the same cluster. We identified 11 dSSRs between PTUs belonging to cluster 1 (representing regions with more open chromatin), 51 dSSRs between PTUs in cluster 2, and 11 dSSRs between PTUs in cluster 3 (indicating regions with more closed chromatin). Statistical analysis was performed using one-way ANOVA with Tukey’s range test as post-hoc (adjusted p-value: 0.0034).

### Supporting Tables

**S1 Table.** Total and mapped reads

**S2 Table.** Raw and processed data of all genomic features classified by their genomic compartment and CDSs per PTU

**S3 Table.**TPM values all PTUs classified by their genomic compartment, number of CDSs per PTU, presence of adjacent ncDNA

**S4 Table.** Genome features with undectable levels of nascent expression (GROexp- GRObck<=0)

**S5 Table.** FPKM values of epigenetic marks and nucleosome occupancy in epimastigote forms from all dSSRs classified by genomic compartment and number of CDSs per PTU

**S6 Table.** Abundance values from 5mC (m) and 5hmC (h) in dSSRs from epimastigote forms classified by genomic compartment and strand

